# Optimizing Parameters for Using the Parallel Auditory Brainstem Response (pABR) to Quickly Estimate Hearing Thresholds

**DOI:** 10.1101/2021.05.13.444069

**Authors:** Melissa J Polonenko, Ross K Maddox

## Abstract

**Objectives:** Timely assessments are critical to providing early intervention and better hearing and spoken language outcomes for children with hearing loss. To facilitate faster diagnostic hearing assessments in infants, the authors developed the parallel auditory brainstem response (pABR), which presents randomly timed trains of tone pips at five frequencies to each ear simultaneously. The pABR yields high-quality waveforms that are similar to the standard, single-frequency serial ABR but in a fraction of the recording time. While well-documented for standard ABRs, it is yet unknown how presentation rate and level interact to affect responses collected in parallel. Furthermore, the stimuli are yet to be calibrated to perceptual thresholds. Therefore, this study aimed to determine the optimal range of parameters for the pABR and to establish the normative stimulus level correction values for the ABR stimuli.

**Design:** Two experiments were completed, each with a group of 20 adults (18 – 35 years old) with normal hearing thresholds (≤ 20 dB HL) from 250 to 8000 Hz. First, pABR electroencephalographic (EEG) responses were recorded for six stimulation rates and two intensities. The changes in component wave V amplitude and latency were analyzed, as well as the time required for all responses to reach a criterion signal-to-noise ratio of 0 dB. Second, behavioral thresholds were measured for pure tones and for the pABR stimuli at each rate to determine the correction factors that relate the stimulus level in dB peSPL to perceptual thresholds in dB nHL.

**Results:** The pABR showed some adaptation with increased stimulation rate. A wide range of rates yielded robust responses in under 15 minutes, but 40 Hz was the optimal singular presentation rate. Extending the analysis window to include later components of the response offered further time-saving advantages for the temporally broader responses to low frequency tone pips. The perceptual thresholds to pABR stimuli changed subtly with rate, giving a relatively similar set of correction factors to convert the level of the pABR stimuli from dB peSPL to dB nHL.

**Conclusions:** The optimal stimulation rate for the pABR is 40 Hz, but using multiple rates may prove useful. Perceptual thresholds that subtly change across rate allow for a testing paradigm that easily transitions between rates, which may be useful for quickly estimating thresholds for different configurations of hearing loss. These optimized parameters facilitate expediency and effectiveness of the pABR to estimate hearing thresholds in a clinical setting.

**Highlights:**

- The pABR yields robust responses across stimulus rates and intensities.
- The optimal rate is 40 Hz, but using multiple rates may prove useful.
- The pABR shows some adaptation with increased stimulation rate.
- Extended analysis windows improve response detection for low stimulus frequencies.
- Behavioral thresholds subtly change across pABR rate, giving similar dB nHL values.

## INTRODUCTION

Early identification of hearing loss and timely intervention is important for promoting typical auditory development and spoken language acquisition (Ching et al. 2014; Cullington et al. 2017; May-Mederake 2012; Moeller 2000; Niparko et al. 2010). Currently the gold standard for estimating hearing thresholds in young infants, and other individuals who do not provide reliable behavioral responses, involves serially measuring electrophysiological auditory brainstem responses (ABRs) to frequency-specific tone pip stimuli presented over a range of intensities to each ear separately (American Academy of Audiology 2012; BC Early Hearing Program 2012; NHS Newborn Hearing Screening Programme 2013; Ontario Ministry of Children, Community and Social Services 2018). These diagnostic ABRs can take a long time, and often yield incomplete information because the child cannot sleep or remain still long enough to obtain the necessary responses. Multiple visits to obtain a complete assessment delays treatment, carries additional costs and risk of attrition, adds stress to the family as they await clinical decisions, and burdens clinician times and resources. To address the time constraints of testing, we recently validated the new parallel ABR (pABR) as a viable method for facilitating faster recording of canonical ABR waveforms than traditional serial methods (Polonenko & Maddox 2019). However, the optimal presentation rates across level for the pABR have yet to be established. In this paper we determined the optimal parameters for using the pABR to estimate hearing thresholds.

The pABR method uses time-saving strategies of simultaneous presentation of stimulus sequences at five frequencies to both ears and randomized stimulus timing sequences. Simultaneous presentation has been a successful tool used for estimating hearing thresholds with the auditory steady-state response (Luts et al. 2006; Sininger et al. 2018; Van Maanen & Stapells 2010). While simultaneous presentation allows for multiple responses to be recorded, randomization allows for unrestricted analysis windows, affording higher stimulation rates and better estimates of the pre-stimulus noise (Burkard et al. 1990; Eysholdt & Schreiner 1982; Polonenko & Maddox 2019; Valderrama et al. 2016; Valderrama et al. 2014; Wang et al. 2013). These in turn provide better estimates of signal-to-noise ratio (SNR), a key metric that dictates testing time. However, the actual SNR gains achieved by higher rates become a trade-off between the reduction in noise and the shrinkage of response amplitudes due to neural adaptation at higher rates (e.g., Burkard et al. 1990; Burkard & Hecox 1983; Chiappa et al. 1979; Don et al. 1977; Jiang et al. 2009).

The optimal range for stimulation rate across intensities is well studied for serial ABR measurement, but the effects of simultaneous stimulation across all frequencies with random timing are not obvious. Parallel presentation across frequency bands and ears may lead to cochlear excitation patterns that differ from the standard single-frequency ABR. Indeed, each frequency may act as a masker for the other frequencies, particularly at higher levels when there is more spread of activation along the cochlea. Preliminary evidence for different interactions comes from the longer latencies and smaller amplitudes of wave V for the pABR than serial ABR, particularly at higher intensities and lower frequencies (Polonenko & Maddox 2019). Potential interactions that depend on both stimulus level and rate may result in optimal stimulus parameters for the pABR that differ from the standard ABR.

Clinical application of the pABR for threshold estimation will also depend on accurate calibration of the pABR stimuli. There are two considerations for calibration due to the short duration of tone pip stimuli. First, transient stimuli such as tone pips are physically calibrated in peak equivalent SPL (dB peSPL) by matching the amplitude of the tone pip to that of a 1000 Hz tone because the time constants of sound level meters are too long to adequately capture the level of the short stimuli (Laukli & Burkard 2015). The amplitudes can be matched by the baseline-to-peak (peSPL) or peak-to-peak (ppeSPL) of the transient (for a discussion of the merits of both approaches see Laukli & Burkard 2015). We chose to calibrate our pABR stimuli in dB peSPL since our stimuli were slightly asymmetric and for the ease of comparing to clicks and converting to other metrics such as peak SPL (pSPL). Second, transient stimuli are then psychoacoustically calibrated into units of dB normal hearing level (dB nHL) because perceptual sensitivity in dB peSPL (or dB SPL for tones) varies by frequency. Due to temporal integration, perceptual thresholds for brief stimuli such as tone pips are elevated compared to tones, and vary by pip duration and stimulation rate (Gorga et al. 1984; Gorga & Thornton 1989; Sharma et al. 2003; Watson & Gengel 1969). Therefore, the correction factors for converting thresholds in dB peSPL to the flattened curve of 0 dB nHL are specific to the tone pip parameters and transducers.

Therefore, in this study we investigated the effects of stimulus level and presentation rate on responses measured using the pABR method, with the goal of determining the optimal stimulus parameters and dB peSPL to dB nHL correction factors before evaluating clinical implementation of the pABR for threshold estimation. We show that the pABR yields responses with good SNR that can be recorded over a wide range of rates in reasonable recording times. Furthermore, we show that extending the analysis window to include additional components of the response offers advantages for response detection in many subjects, particularly for the broader responses to low frequency tone pips. Correction factors for perceptual thresholds were relatively similar (within 3.5–6 dB) across stimulation rates.

## MATERIALS AND METHODS

We completed two experiments. First, pABR electroencephalographic (EEG) responses were recorded to different rates and intensities. Second, behavioral thresholds were measured for pure tones and for the pABR stimuli at each rate to determine the correction factors that relate the stimulus level in dB peSPL to perceptual thresholds in dB nHL.

### Subjects

EEG responses and psychoacoustic perceptual thresholds were collected in two separate experiments, each with a different set of 20 adults (13 females, 6 males, 1 non-identifying person for experiment 1; and 11 females, 9 males for experiment 2). There was an additional subject recruited for perceptual threshold testing because one subject was excluded due to unreliable, sporadic thresholds from repeatedly falling asleep during behavioral testing. All subjects gave informed consent before participating in the experiments, which were done under protocols approved by the University of Rochester Institutional Review Board (#3866). The mean ± SD (range) age was 22.5 ± 4.2 (18–35) years for EEG testing, and 21.8 ± 3.4 (18–33) years for behavioral testing.

Normal hearing thresholds, defined as ≤ 20 dB HL, were confirmed at octave frequencies from 250–8000 Hz. Tympanometry and otoscopy confirmed normal middle and outer ear function. Distortion product emission (DPOAE) testing confirmed normal outer hair cell function from 1–4 kHz, barring 2 subjects who had a DPOAE but poor SNR at 2 kHz in one of their ears. Most subjects also had DPOAEs at 500 Hz, but the noise floor was high and the DPOAE did not reach the 6 dB SNR criteria for one ear in 2 subjects and for both ears in 4 subjects.

### pABR stimuli

Details of the pABR stimulus construction and method can be found in Polonenko and Maddox (2019). Briefly, stimuli to each ear comprised summed independent, randomly timed trains of Blackman windowed 5-cycle cosine tone pips centered at octave frequencies from 500 to 8000 Hz. Individual tone pips had durations of 10, 5, 2.5, 1.25 and 0.625 ms for the frequencies 500, 1000, 2000, 4000 and 8000 Hz respectively. Thirty unique 1 s stereo epochs were created to ensure sufficient statistical independence between the pseudorandom Poisson processes controlling timing of the tone pip trains (Maddox & Lee 2018; Polonenko & Maddox 2019). To create these tone pip trains, unit-height impulses were randomly inserted across 1 s and convolved with the tone pip. The number of impulses corresponded to the stimulus presentation rate. Polarity was randomly set to ±1 so that half the tone pips were condensation and the other half rarefaction. For the EEG experiment, an inverted version of each of the 30 unique epochs was presented in sequence to counter-phase the stimuli to help mitigate stimulus artifact (i.e., Epoch A^+^, A^−^, B^+^, B^−^, etc, where A and B represent independent epochs and the superscript sign denotes the phase). For the behavioral experiment, a token from these 30 unique epochs was randomly chosen with replacement for each presentation of the stimulus.

The tone pip stimuli were calibrated to 80 dB peak-equivalent SPL (peSPL) by matching the amplitudes of the peak tone pip cosine component to the amplitude of a 1000 Hz sinusoid tone that read 80 dB SPL on a sound level meter (2240, Bruel & Kjaer) when played through an insert earphone (ER-2, Etymotic Research) attached to a 2-cc coupler (RA0038, G.R.A.S.). The other stimulus levels (*L*) in dB peSPL were obtained by multiplying the reference-level tone pip by 10^(*L* − 80) / 20^. For the behavioral experiment, pure tones were created with amplitudes that were also calibrated to the same 80 dB SPL tone.

Stimuli were created at a sampling rate of 48 kHz and presented through ER-2 insert earphones connected to a sound card (Babyface Pro, RME, Haimhausen, Germany). For the EEG, the sound card was also connected to a headphone amplifier (HB7, Tucker Davis Techologies, Alachua, FL, USA), which sent an inverted stimulus to a second “dummy” set of earphones that had a blocked tube and was taped in the same orientation to the stimulus earphones. This setup further mitigated stimulus artifact by cancelling electromagnetic fields close to the transducers. The earphones were also hung from the ceiling by magnets to allow as much distance as possible between the transducers and EEG electrodes. Stimulus presentation was controlled by a python script using publicly available software at https://github.com/LABSN/expyfun (Larson et al. 2014). The sound card’s optical digital out was also used to send digital signals that precisely marked the beginning of each 1 s epoch, which were then converted to trigger pulses by a custom trigger box (modified from a design by the National Acoustic Laboratories, Sydney, NSW, Australia). These triggers were then sent to the EEG system in order to synchronize responses with stimuli.

### EEG experiment

#### Stimulus conditions and EEG recording

We recorded responses in both ears to pABR stimulation with average presentation rates of 20, 40, 60, 80, 100, and 120 Hz, each at intensities of 51 and 81 dB peSPL. For a single 2-hour recording session, this afforded 10 minutes for each of the 12 conditions, resulting in averaged responses comprised of 12,000 (20 Hz rate) to 72,000 (120 Hz rate) repetitions. Conditions were interleaved to prevent changes in impedance, subject state, or EEG noise from affecting one condition more than the others.

Two-channel scalp potentials were recorded with BrainVision’s PyCorder software, using passive Ag/AgCl electrodes connected to two differential preamplifiers (BrainVision LLC, Greenboro, SC). In standard 10–20 coordinates, the non-inverting (positive) electrode was positioned at FCz (just anterior of the vertex), the inverting (negative) electrodes at A1 and A2 (left and right earlobes), and the ground electrode at FPz (frontal pole). The non-inverting and ground electrodes were plugged into y-connectors that split between the two differential pre-amplifiers. Data were sampled at 10 kHz and high-pass filtered at 0.1 Hz. Triggers marked the beginning of each epoch rather than each individual tone pip stimulus for two reasons: 1) to avoid trigger overlap due to random stimulation of 10 tone pips (5 for each ear), and 2) to efficiently analyze 1 s blocks of data in the frequency domain, which is mathematically equivalent to – but faster than – averaging responses to each individual tone pip. Participants reclined in a darkened sound treated audiometric booth during testing, and were encouraged to rest.

#### pABR response calculation

Further details can be found in Polonenko and Maddox (2019) but a brief description is provided below. Using the mne-python package (Gramfort et al. 2013), raw EEG was filtered offline from 30–2000 Hz using a first order causal Butterworth filter, and then notch-filtered at odd multiples of 60 Hz up to 2500 Hz to remove electrical line noise.

As mentioned above, triggers denoted the beginning of each 1 s epoch. This epoch, along with 500 ms before and after it was extracted for each trigger, giving 2 s of EEG data, denoted as *y*. This same EEG was used to derive responses for each of the 10 tone pips for that epoch. For each tone pip of frequency, *f*, and ear, *e*, we took the rectified impulse sequence used to create the tone pip train – a unit-height impulse at the onset of each tone pip in the 1 s epoch – and zero-padded it with 500 ms before and after to give a 2 s impulse train with all the impulses in the center 1 s, which was denoted *x_f,e_*. The response, *w_f,e_* was then calculated as the circular cross-correlation of the 2 s EEG and the 2 s zero-padded rectified pulse train, performed in the frequency domain as *w_f,e_* = 1/*n F*^−1^ {*F*{*x_f,e_*}* *F*{*y*}} where *F* denotes the fast Fourier transform, *F*^−1^ its inverse, * denotes complex conjugation, and *n* the number of impulses in a sequence (e.g., 40 for the 40 Hz stimulation rate). Due to the circular nature of the cross-correlation, the time interval [0, 500] ms was at the beginning of *w_f,e_* and [−500, 0) ms at the end.

Concatenating these two time intervals (i.e., discarding the middle 1 s of *w_f,e_*) gave the final response from [−500, 500] ms, where 0 ms denotes the onset of the tone pip. This was repeated for each of the 10 tone pips and for each of the two EEG channels for every epoch.

Average responses for each condition (level, rate, channel, ear, and tone pip frequency) were calculated by first weighting each epoch by the inverse variance of its pre-stimulus baseline from −480 to −20 ms relative to the summed pre-stimulus inverse variances of all epochs for that condition, and then summing across the weighted epochs. This method resembles Bayesian averaging (Elberling & Wahlgreen 1985), but leverages the much longer 500 ms pre-stimulus baseline afforded by random stimulus timing. Using this averaging method, very noisy epochs contribute much less to the grand average by assigning a weight close to zero. This avoids the need for rejecting epochs based on threshold criteria. Responses were also averaged across channels to reduce noise because we were not interested in ipsilateral versus contralateral differences for this paper. However, it would be easy to keep the two channels separate for clinical applications.

#### EEG data analysis

The primary objective of this paper was to determine the optimal rate for quickly obtaining waveforms for all 10 tone pips. To achieve this, we: 1) compared the wave V latency (ms) and amplitude (μV) across conditions using linear mixed effect regression with rate and intensity fixed effects and a random intercept per subject; 2) quantified the response SNRs; and 3) estimated the recording time required for all responses in a rate-intensity combination to achieve 0 dB SNR. We chose a threshold of 0 dB SNR based on when waveforms became clearly identifiable and what we have done previously (Maddox & Lee 2018; Polonenko & Maddox 2021; Polonenko & Maddox 2019). Of course, other dB SNR thresholds would change the estimated times to reach this criterion, but in a proportional way for each condition. Two analysis time windows were tested to determine SNR: a 10 ms window to include wave V of the ABR, and a 30 ms extended window to include wave V of the ABR and early components of the middle latency response (MLR).

The dB SNR of each averaged response (i.e., after 600 epochs, or 10 minutes) was estimated according to the formula: SNR_600_ = 10log_10_[(σ^2^_S+N_ − σ^2^_N_) / σ^2^_N_] where σ^2^_N_ was the variance of the noise calculated as the mean variance over 10 ms (ABR wave V window) or 30 ms (ABR/MLR extended window) intervals from −480 to −20 ms, and σ^2^_S+N_ was the variance of the signal and noise calculated as the variance in the respective 10 ms or 30 ms latency range starting at a lag that captures wave V for each tone pip’s frequency: 10.5, 7.5, 6.5, 5.0, and 5.0 ms for 500, 1000, 2000, 4000 and 8000 Hz respectively (Polonenko & Maddox 2019; Stapells 2010). Then we standardized the SNR to a 1 minute (60 s) run: SNR_60_ = SN_600_ + 10log_10_(60 / 600). From SNR_60_ we estimated the time-to-0 dB SNR as 60 × 10^−SNR60 / 10^. This time was calculated for each tone pip, but the overall acquisition time for a condition is based on the slowest waveform, and as such, was calculated as the maximum time-to-0 dB of the 10 simultaneously acquired waveforms. Cumulative density functions were computed across subjects of time-to-0 dB for each tone pip in order to determine the optimal presentation rate – the rate at which 90% of subjects reached an SNR ≥ 0 dB for all 10 tone pips in the shortest recording time.

The linear mixed effects regressions and their power analyses were performed using the lmer4, lmerTest and simR packages in R (Bates et al. 2015; Green & MacLeod 2016; Kuznetsova et al. 2017; R Core Team 2020). To calculate power, the likelihood ratio test was performed on 1000 Monte Carlo permutations of the response variables based on the fitted model.

### Perceptual thresholds experiment

#### Stimulus conditions and psychoacoustics parameters

The 1 s stimulus tokens used for EEG were also used for determining perceptual thresholds for dB nHL correction factors. However, the tone pip trains were not presented in parallel but serial to determine the threshold for each tone pip frequency. In addition to the tone pip stimuli, we measured thresholds for 1 s cosine pure tones at each frequency, which were calibrated using the same 1000 Hz tone at 80 dB SPL that was used to calibrate the pABR tone pips. The tones had raised cosine window edges set to 35 ms, which was within the American National Standards Institute (ANSI) S3.6-2010 rise/fall standards and matched that of our audiometer used for hearing screening.

During the behavioral task, subjects sat in a sound treated audiometric booth, looked at a dark computer monitor screen with a central white fixation dot, and responded with keyboard presses when they heard the stimulus. Stimulus presentation time was jittered by a random number between 1 to 4 s. Perceptual thresholds were measured using an automated tracker based on the modified Hughson Westlake method (Carhart & Jerger 1959), with a 5 dB base step size in a 2-down/1-up paradigm and a starting level of 40 dB (pe)SPL. If there was no response for the starting level, then the level was increased in 20 dB steps until the subject responded or a maximum level of 85 dB (pe)SPL was reached. Threshold was defined as the level at which 2 out of 3 correct responses were given when the level of the stimulus was ascending. All 70 threshold tracks were randomly presented over the course of the experiment (2 ears × 5 frequencies × 7 rates, with 0 Hz rate representing tones). For each presentation of a pABR tone pip, a random token was chosen with replacement from the thirty unique 1 s tokens. Breaks of at least 15 s were provided after every 4 threshold tracks, and subjects chose when to continue after a break if they needed longer than 15 s to rest.

### Data analysis

Attentional state drifted in some subjects resulting in a few spuriously high thresholds. Thresholds were considered outliers and removed when the threshold to the pABR stimulus was >40 dB above that for the pure tone, and the track was confirmed to be poor and reflecting dozing (i.e., the intensity increased to 85 dB peSPL with no response). Of 1,400 thresholds, 23 (1.6%) were removed and there was never more than 3 of 40 data points removed for a frequency-rate condition.

The reference values in dB nHL were calculated similarly to what has been done before (Gorga et al. 1993; Sharma et al. 2003; Stapells & Oates 1997). The thresholds to pure tones were subtracted from the thresholds to the brief pABR stimuli to correct for temporal integration and the subject’s own pure tone thresholds. These corrections were modeled using linear mixed effects regression with fixed effects of rate, logged frequency, and their 2-way interaction, as well as a random intercept for subject-ear. Then the corrections derived from the model for each rate-frequency condition were converted to reference values to give 0 dB nHL by adding them to the HA-1 coupler reference equivalent threshold in SPL (RETSPL) for the ER-2 earphones (i.e., the conversion of SPL to audiometric zero).

## RESULTS

### The pABR yields canonical brainstem responses that characteristically show adaptation at higher rates

We recorded the pABR over a range of stimulation rates in 20 Hz steps and at two intensities. Figure 1 shows the grand average waveforms for each of these conditions, and for each ear and tone pip frequency. Overall, morphology of the pABR responses resembled the canonical responses from traditional ABR methods, with lower frequency responses exhibiting a broader wave V than the higher frequencies. Although wave V is the primary focus of methods for estimating hearing thresholds, Figure 1 also shows that additional ABR waves I and III were clearly visible in the higher frequency responses across several presentation rates, especially at the higher intensity of 81 dB peSPL.

**Figure 1.**
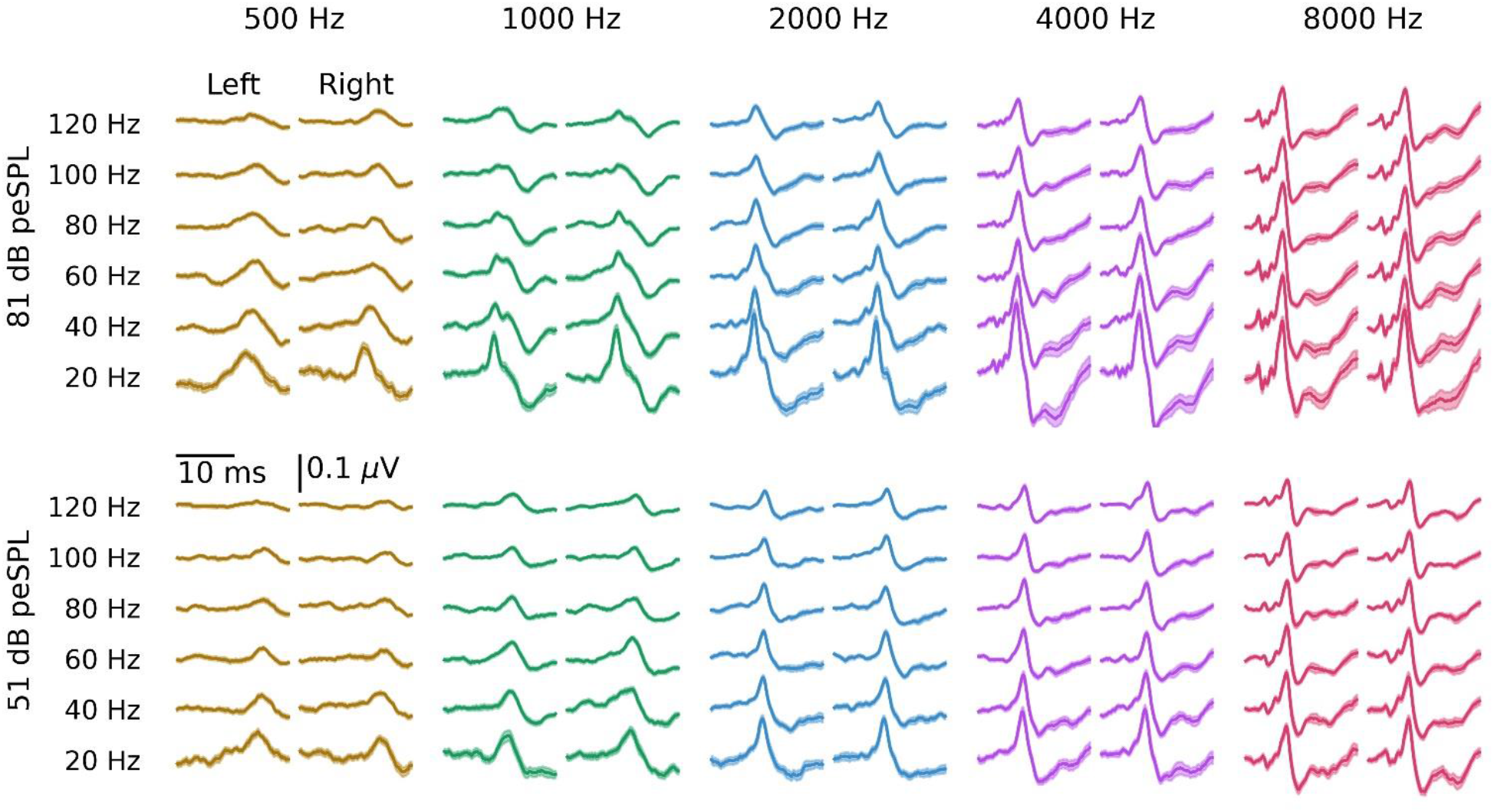
Grand average waveforms across stimulation rates and tone pip frequencies, plotted for both the left and right ears and for a high (top) and low (bottom) intensity. Areas show ± 1 SEM, computed across subjects. All responses are plotted over the interval 0 to 20 ms.

Waveforms were visually inspected and quantified for the presence, amplitude and latency of wave V. Amplitude was defined as the peak to following trough. All 2,400 responses were quantified by a trained audiologist (MJP), and the other author (RKM) quantified a subset of 720 responses (30%) from a random selection of 6 participants. Wave V was manually quantified to mimic what is done in the clinic and to avoid noise spuriously being identified as the peak, especially in the broader low frequency responses. The intraclass correlation coefficient for absolute agreement (ICC3) was ≥0.81 for each frequency and measure (the lowest two ICC3 95% confidence intervals were 0.76–0.85 and 0.89–0.94 for 500 Hz latency and 1000 Hz amplitude respectively, all others were ≥0.94), indicating good agreement for chosen wave V peaks. In 13 waveforms there was not a clear wave V and following trough compared to the recording noise. These absent responses were for 500 Hz at the two highest stimulation rates of 100 Hz (n = 6) and 120 Hz (n = 7), and mostly for 51 dB peSPL (3 / 13 at 81 dB peSPL). For further analyses, the latencies of missing waves were removed and the amplitudes considered to be zero.

We modeled wave V latency (Figure 2A) and amplitude (Figure 2B) using linear mixed effects models, with a random intercept for each subject and fixed factors of rate (for amplitude measures only, rate was log transformed due to the non-linear relationship), intensity, ear, frequency in log units, and gender. We included the ear–frequency interaction, as well as all 2-way and 3-way interactions between rate, intensity, and frequency. There was one subject who did not identify as male or female and could not be included in the full model due to insufficient numbers for a third gender category. Details of the full statistical models for latency and amplitude are given in Supplemental Digital Content 1 (see Table 1A and 1B respectively). There was not a significant effect of gender for latency (0.22 ± 0.16 ms, *t*(19) = 1.38, *p* = 0.184, power = 0.29 [95% confidence interval: 0.26–0.32]), but there was a trend that responses from female subjects were slightly larger (0.025 ± 0.012 μV [95% confidence interval = −0.001–0.049 μV], *t*(19) = 2.05, *p* = 0.054, power = 0.55 [95% confidence interval = 0.52–0.58]). To include all 20 subjects, we confirmed that the significant effects in the full model were maintained in models excluding the non-significant fixed effect of gender, and the details of these statistical models are found in Table 1. Wave V latency showed a difference between ears (*p* = 0.003) but also a significant ear–frequency interaction (*p* = 0.008), indicating that responses for the right ear were faster for lower frequencies (mean ± SEM difference: 0.28 ± 0.07 ms for 500 Hz) but similar for higher frequencies (0.02 ± 0.03 ms difference for 8000 Hz). Consistent with our previous study (Polonenko & Maddox 2019), there were also significant effects of intensity, frequency, and a significant intensity–frequency interaction (all *p* < 0.01), indicating that latency decreased with increasing frequency and increasing intensity, and the effect of intensity was greater at lower frequencies. The slight increase in latency with increasing rate was not significant (*p* = 0.875), and there was no significant rate–intensity, rate–frequency, or rate– intensity–frequency interactions (all *p* > 0.109). Unlike latency, wave V amplitude showed no effect of ear or an ear–frequency interaction (both *p* > 0.118). Also consistent with our previous study (Polonenko & Maddox 2019), wave V amplitude increased with increasing intensity at a greater rate for higher frequencies (intensity–frequency interaction, *p* < 0.001). Here, we also showed that amplitude decreases with increasing rate to a greater extent at higher intensities and higher frequencies (rate–intensity–frequency interaction, *p* = 0.010). There was an exception for 8000 Hz, which showed similar or smaller amplitudes than 4000 Hz for the lower rates.

**TABLE 1.**
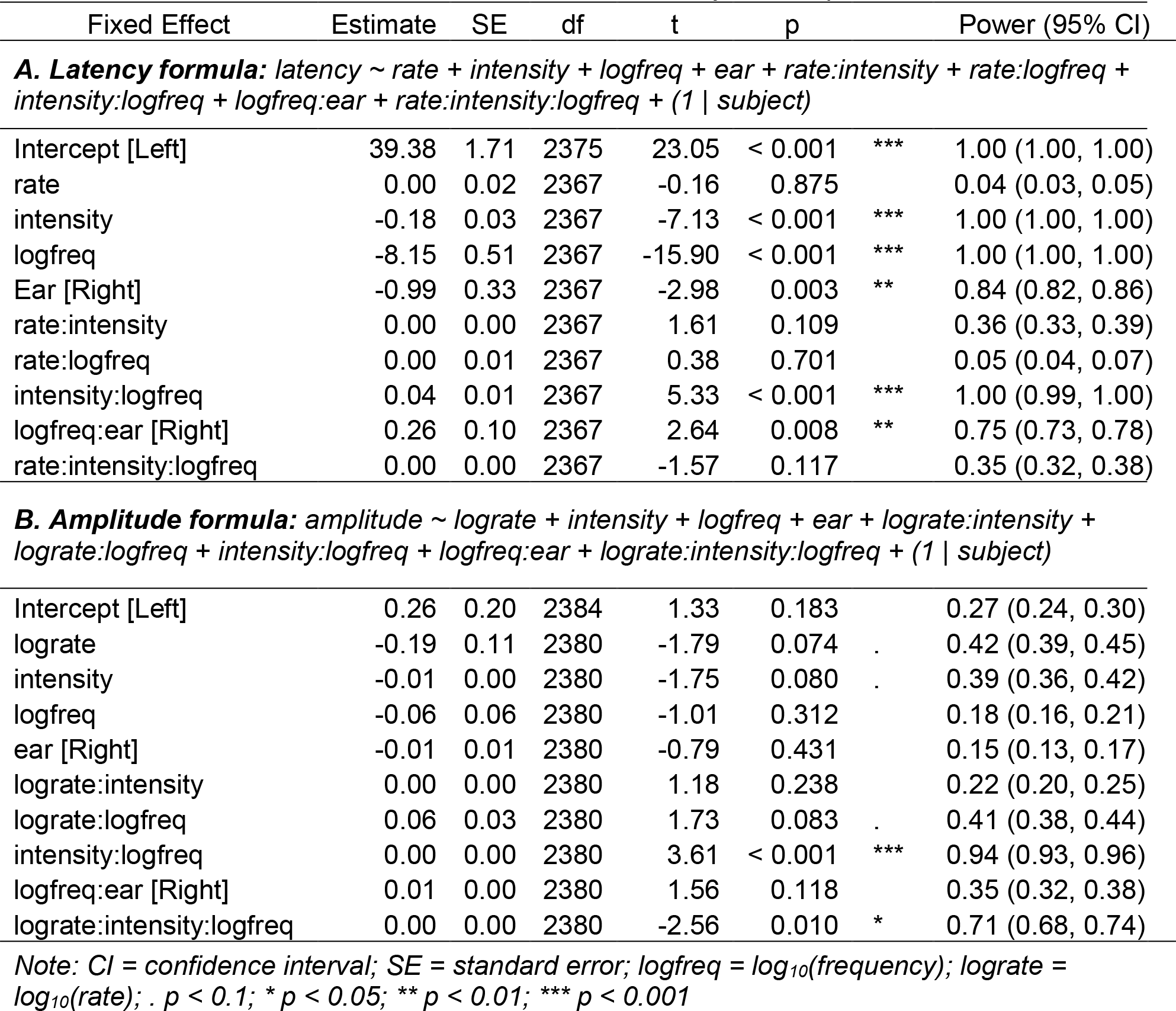
Linear Mixed Effects Models for Wave V Latency and Amplitude

**Figure 2.**
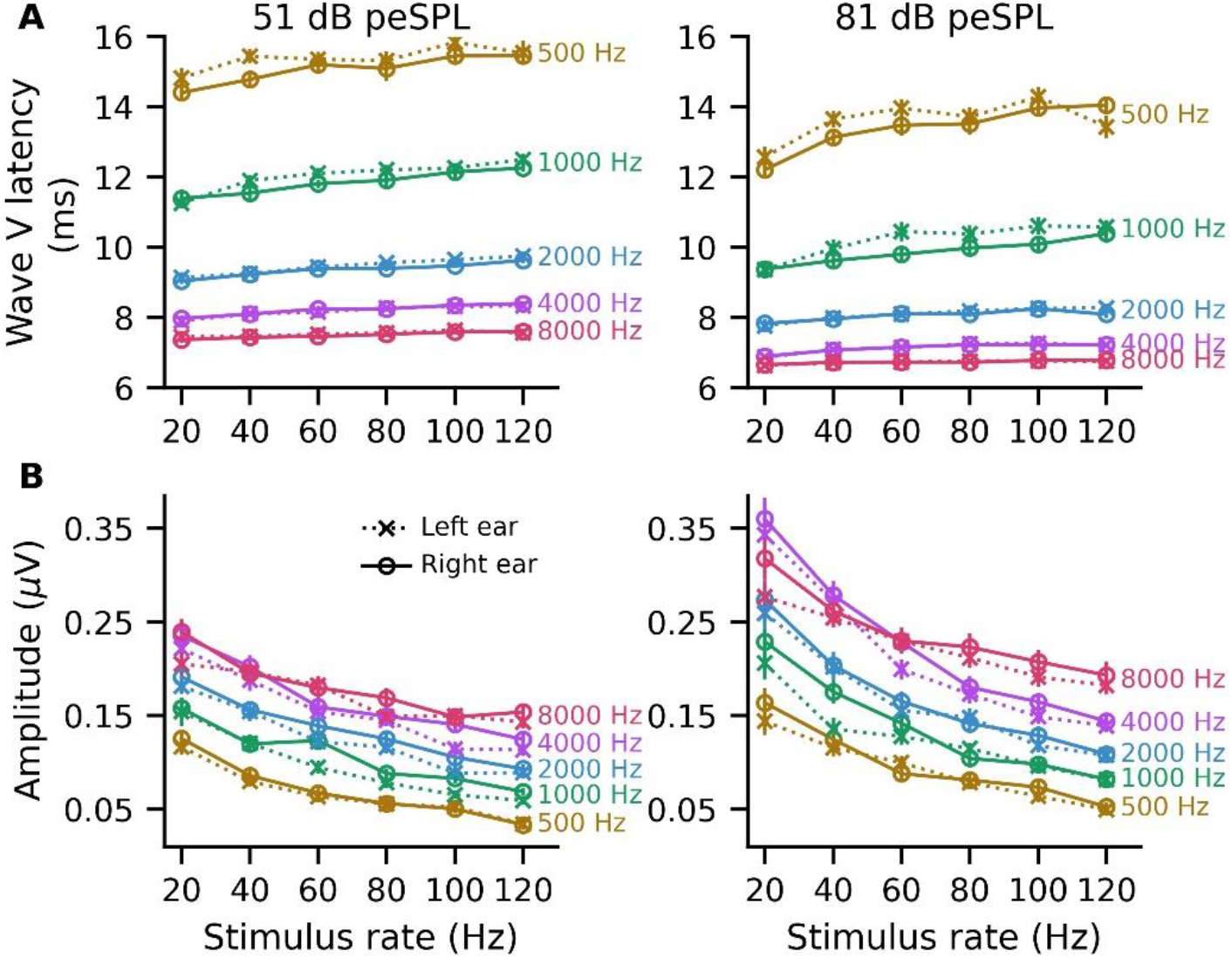
Mean wave V latency (A) and amplitude (B) as a function of stimulation rate at a low (left) and high (right) intensity. Stimulus frequency is indicated beside each line. Error bars (where large enough to be seen) indicate ± 1 SEM. Crosses joined by dotted lines indicate left ear responses and circles joined by solid lines indicate right ear responses.

### Acquisition times are fastest for a stimulation rate of 40 Hz

Next, we explored how the changes in amplitude with rate affect the time required for responses to reach ≥ 0 dB SNR using a 10 ms analysis window. We extrapolated to a maximum of 20 minutes, which was twice our recording time but represented the minimum time required for serial collection of each response based on our previous study (Polonenko & Maddox 2019). To determine the optimal pABR rate for the majority of subjects, the cumulative proportion of subjects was computed as a function of recording time. The cumulative density functions (CDF) for the 40 Hz rate are shown in Figure 3A, and the CDFs for all rates are provided in Supplemental Digital Content 2 (see Figure 1). Figure 3B shows the time to 0 dB SNR for 90% and 50% (median time) of responses for each rate, which were taken from the CDFs.

**Figure 3.**
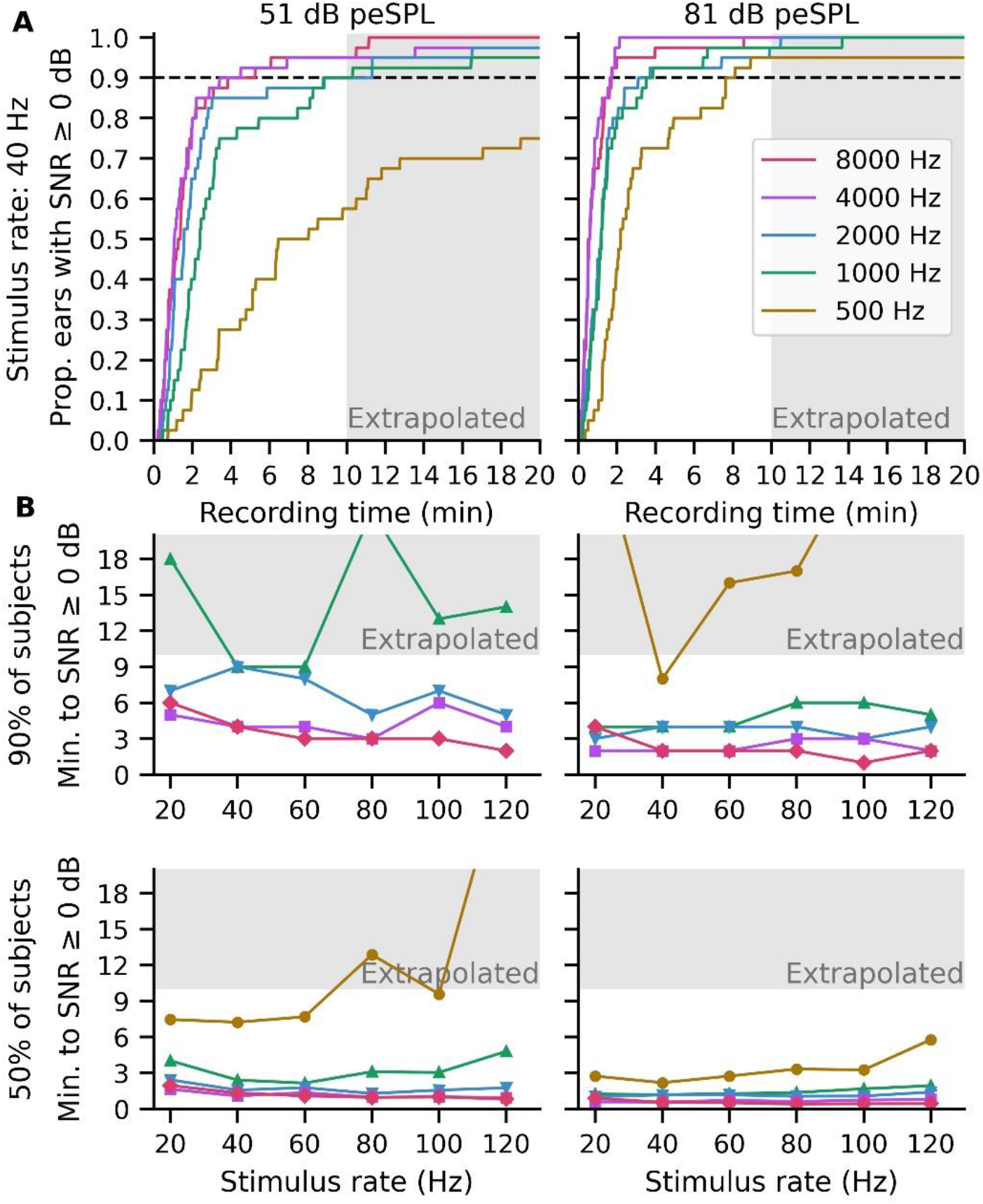
Acquisition times for waveforms to reach ≥ 0 dB SNR for a low (left) and high (right) presentation level for each tone pip frequency (indicated by line color). (A) The cumulative density function (CDF) is shown for the 40 Hz stimulation rate. From the CDF for each stimulation rate, the time for waveforms to reach ≥ 0 dB SNR in 90% and 50% of subjects (median) were calculated and displayed in (B). Shaded areas represent time estimations extrapolated past the 10-minute recording time.

Acquisition times were faster for the higher tone-pip frequencies and lower stimulation rates. Half the subjects had all waveforms within 9.6 minutes for 51 dB peSPL and 3.5 minutes for 81 dB peSPL, except for the 120 Hz rate at 81 dB peSPL (5.8 minutes) and for the 80 Hz and 120 Hz rates at 51 dB peSPL (12.9 and > 20 minutes respectively). In fact, for a 40 Hz stimulation rate the median time was 7.3 and 2.2 minutes for 51 and 81 dB peSPL. However, the acquisition time necessary for most subjects was limited by the 500 Hz tone pip – the broadest and lowest-amplitude response. The estimated time for ≥ 90% of subjects to reach 0 dB SNR for 2000–8000 Hz was ≤ 9 and 4 minutes for 51 and 81 dB peSPL respectively. But for 500 Hz the acquisition time – and thus, the total time for the pABR – was > 20 minutes for all rates at the lower intensity, and 7.7 minutes for 40 Hz but >15 minutes for the other rates at the higher intensity. At the end of the 10-minute recording session, 48–72% of the 500 Hz responses reached criterion at the lower intensity and 60–92% reached criterion at the higher intensity. Based on the 50% and 90% metrics, a 40 Hz stimulation rate appears optimal for ensuring the timeliest robust responses from most subjects, although rates between 20–60 Hz required similar times for the mid and higher frequencies.

### Extended analysis windows afforded by random timing improves SNR and saves time for low frequencies in some cases

The random timing of the pABR allows for extending the analysis window to view more components of the evoked response. This advantage may improve SNR and acquisition estimates for the broader low frequency responses, which are present but vary less over 10 ms than the high frequency responses. Figure 4 shows the same responses from Figure 1 with the time window extended from 20 to 40 ms, which allows an analysis window of 30 ms for each tone pip frequency. With this extended window, the negative trough following wave V is now visible for the lower frequency responses (along with the higher frequency responses, which were visible with the shorter window), as well as the early components of middle latency response (MLR).

**Figure 4.**
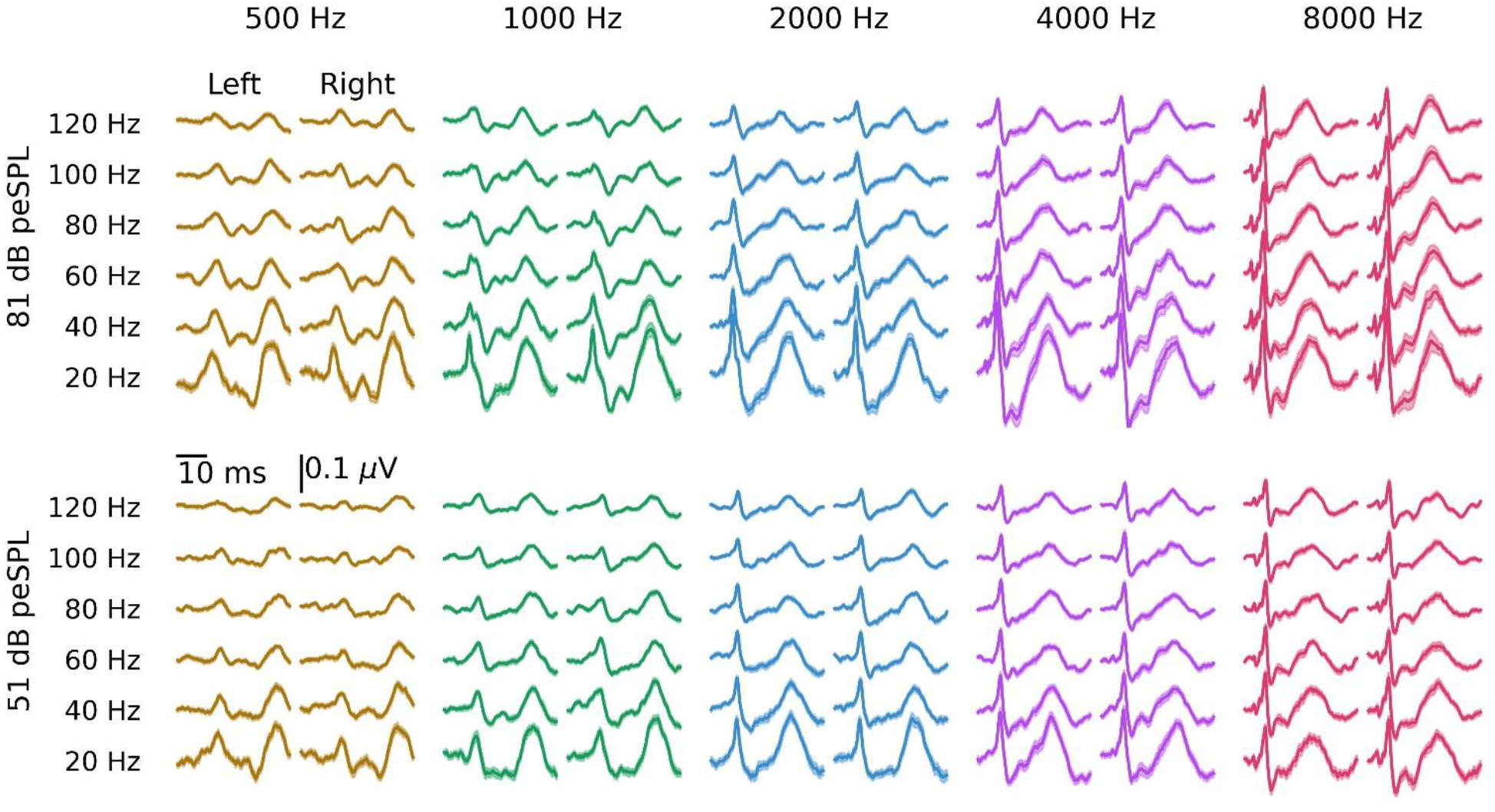
Grand average waveforms across stimulation rates and tone pip frequencies, plotted for both the left and right ears and for a high (top) and low (bottom) intensity. Areas show ± 1 SEM across subjects. All responses are plotted over the interval 0 to 40 ms.

Figure 5 compares the acquisition times for the low frequency tone-pips at low stimulation rates when using a 30 ms versus 10 ms window to estimate SNR. We focused on 20 and 40 Hz because these stimulation rates showed the best recording times with the 30 ms analysis window, as well as the greatest response variation over the extended window for 500 and 1000 Hz tone pips. The CDFs for all rates are provided in Supplemental Digital Content 2 (see Figure 2). For many subjects the 30 ms window provided similar or better estimates of SNR and acquisition times, especially for the 500 Hz tone pip. In Figure 5A, the CDFs for the 20 Hz stimulation rate began with a similar trajectory, but grew faster for the 30 ms window (i.e., diverged from the curve for the 10 ms window) for recording times > 4 minutes (at ~35% of ears) for 500 Hz and > 6 minutes (at ~70% ears) for 1000 Hz at 51 dB peSPL, and after ~5.5 minutes for 500 Hz (at ~72% of ears) at 81 dB peSPL. The 1000 Hz CDF for the 10 ms window was higher than that for the 30 ms window at all times for 81 dB peSPL. This means that the 30 ms analysis window provided a time benefit for the ~65% of 500 Hz waveforms and ~30% of 1000 Hz waveforms that took longer than 4–6 minutes to reach criterion SNR at the lower intensity level, but the 10 ms window was adequate for the quicker waveforms (i.e., < 4–6 minutes) and for most of the 1000 Hz tone pip waveforms at the higher intensity.

**Figure 5.**
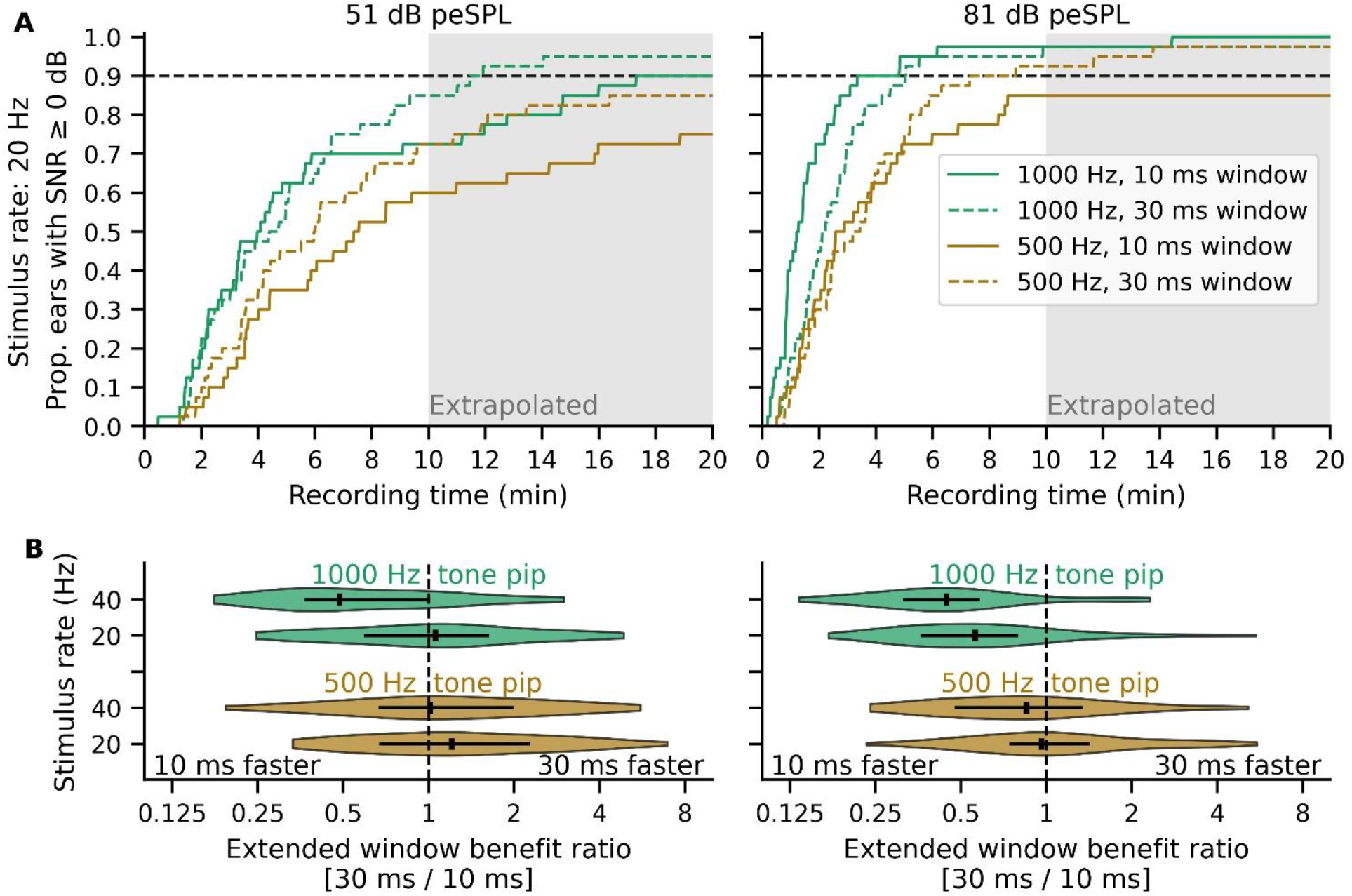
Extending the analysis window from 10 to 30 ms can improve response dB SNR and acquisition time for low frequency tone pips. (A) An example cumulative density function (CDF) is shown for the 20 Hz stimulation rate. Recording time required for waveforms to reach 0 dB SNR with a low (left) and high (right) presentation level for 500 and 1000 Hz tone pips when using an extended analysis window of 30 ms. Stimulus frequency is indicated by line color. (B) The distribution of extended window benefit ratios for using a 30 versus 10 ms analysis window. The horizontal black line indicates the interquartile range and the vertical solid black line denotes the median ratio. Ratios are on a log2 scale, with ratios > 1 indicating a benefit to the 30 ms versus 10 ms window. The 10 ms window includes wave V of the ABR, and the extended 30 ms window also includes components of the middle latency response. The dashed line represents similar times to reach 0 dB SNR for both analysis windows.

Consistent with the CDFs, the distribution of extended window benefit ratios in Figure 5B suggest that the 30 ms analysis window provides a time benefit for the slower set of waveforms, with significant improvements for some subjects. The greatest extended window benefit ratios for a 30 ms window occurred for 500 Hz at 51 dB peSPL. Half the subjects had speedup ratios ≥ 1, and for the “slowest 25%” of subjects (i.e. 75^th^ percentile and higher), the 30 ms window gave extended window benefit ratios of 2–7, corresponding to 5.7–16.1 minutes saved by using a 30 ms window. At this intensity, the 75^th^ percentile extended window benefit ratios were < 1.6 (up to ~3 minutes saved) for 1000 Hz, but there were also some cases that benefited from the 30 ms window by a ratio of 3-5, or up to 13–15 minutes saved. At the higher intensity of 81 dB peSPL, there were faster or similar recording times for the 10 ms window in at least half the subjects, with the median extended window benefit ratios ≤ 1 for both stimulus rates and tone pip frequencies. However, the 30 ms window benefited about half the subjects for 500 Hz, and there were significant improvements for some subjects for both 500 Hz and 1000 Hz. The 75^th^ percentile extended window benefit ratios were < 1 for 1000 Hz and <1.4 for 500 Hz (up to 1.2 minutes saved by the 30 ms window). Again, even for the high intensity, there were some extreme cases for whom the 30 ms provided an extended window benefit ratio up to 5.5, corresponding to a maximum of 5 to 16.5 minutes saved. To succinctly summarize the above, having the option to use a 30 ms window will speed the exam for some patients.

### Behavioral thresholds for pABR stimuli subtly change with rate

Having examined the effects of stimulation rate on the pABR responses and acquisition times, we next compared perceptual thresholds to the pABR stimuli with those of pure tones to obtain normative values. Figure 6A shows the perceptual thresholds in dB SPL for the pure tones and dB peSPL for the pABR stimuli. Thresholds to the pure tones varied between −10 and 10 dB SPL and were elevated for the pABR stimuli, as expected due to temporal integration of brief stimuli. Thresholds for the pABR stimuli subtly decreased with increasing stimulation rate for each tone pip frequency. This pattern was similar for each subject-ear, as indicated by the light colored lines. The difference between the thresholds to pABR stimuli and pure tones ranged between 0 and 40 dB but were on average between 10 and 21 dB depending on rate and tone pip frequency (Figure 6B).

**Figure 6.**
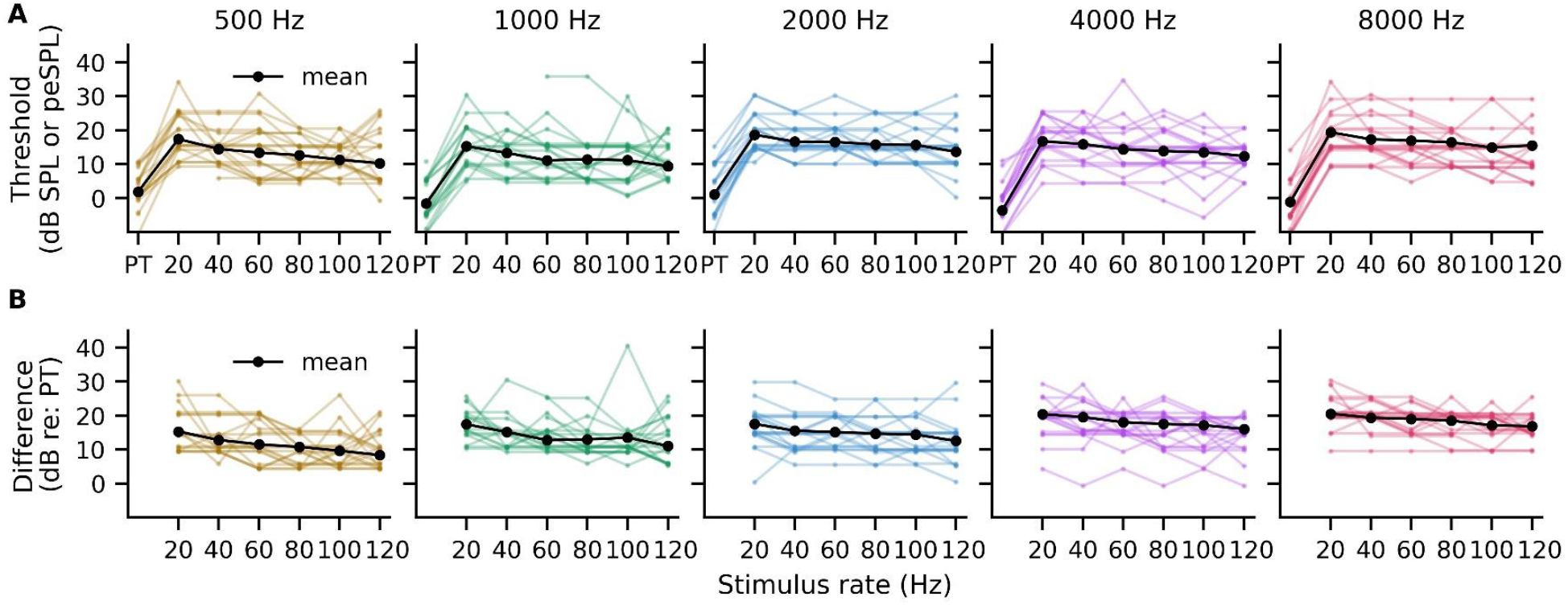
Perceptual thresholds show subtle changes across different stimulation rates. (A) Perception thresholds for pure tones (PT, in dB SPL) and the pABR stimuli (in dB peSPL) with different rates are shown for individuals (color) and the group mean (black). (B) The difference between thresholds to the pABR and pure tone stimuli. Individual colored lines are given a slight random vertical offset between ±1 dB to make the individual data with similar thresholds easier to see.

Next, the normative values for 0 dB nHL were calculated. First, the difference in thresholds, or correction factors, for the pABR stimuli (from Figure 6B) were modeled using linear mixed effects regression. Details of the statistical model are provided in Table 2. As shown in Figure 7A, the threshold difference significantly decreased with increasing stimulation rate (*p* = 0.004) and increased with increasing frequency (*p* < 0.001), but there was not a significant interaction between rate and frequency (*p* = 0.085). The mean difference in correction factor between stimulation rates of 20 and 120 Hz ranged from 3.5 dB for the 500 Hz tone pip to 6.0 dB for the 8000 Hz tone pip. Second, the reference-equivalent thresholds in SPL (RETSPLs) for our ER-2 insert earphones were added to the correction factors to give the normative values for 0 dB nHL (Figure 7B). The values from Figure 7 are also provided in Table 3 for ease of reference.

**TABLE 2.** Linear Mixed Effects Model for pABR Correction Values Model formula: correction ~ rate + logfreq + rate:logfreq + (1 | subject-ear)

| Fixed Effect | Estimate | SE | df | t | p |  | Power (95% CI) |
| --- | --- | --- | --- | --- | --- | --- | --- |
| Intercept | 3.01 | 3.20 | 1162 | 0.94 | 0.348 |  | 0.14 (0.12, 0.16) |
| rate | -0.12 | 0.04 | 1138 | -2.87 | 0.004 | ** | 0.80 (0.77, 0.82) |
| logfreq | 4.74 | 0.96 | 1138 | 4.97 | < 0.001 | *** | 1.00 (0.99, 1.00) |
| rate:logfreq | 0.02 | 0.01 | 1138 | 1.73 | 0.085 |  | 0.39 (0.36, 0.43) |

Note: CI = confidence interval; SE = standard error; $\text{logfreq} = \log_{10}(\text{frequency})$ ;
correction values = threshold differences for pABR tone pip versus pure tone; \* $p < 0.05$ ;
\*\* $p < 0.01$ ; \*\*\* $p < 0.001$

**Table 3.** Correction and normative values for converting pABR dB peSPL to 0 dB nHL

| Tone pip | Stimulus rate |  |  |  |  |  |
| --- | --- | --- | --- | --- | --- | --- |
|  | 20 Hz | 40 Hz | 60 Hz | 80 Hz | 100 Hz | 120 Hz |
| *Model-estimated correction factors (dB)* |  |  |  |  |  |  |
| 500 Hz | 14.6 | 13.4 | 12.2 | 11.0 | 9.8 | 8.6 |
| 1000 Hz | 16.2 | 15.1 | 14.0 | 12.9 | 11.9 | 10.8 |
| 2000 Hz | 17.7 | 16.8 | 15.8 | 14.9 | 13.9 | 13.0 |
| 4000 Hz | 19.3 | 18.5 | 17.6 | 16.8 | 16.0 | 15.2 |
| 8000 Hz | 20.8 | 20.1 | 19.4 | 18.8 | 18.1 | 17.4 |
| *ER-2 normative values for 0 dB nHL (dB)<sup>1</sup>* |  |  |  |  |  |  |
| 500 Hz | 20.6 | 19.4 | 18.2 | 17.0 | 15.8 | 14.6 |
| 1000 Hz | 16.2 | 15.1 | 14.0 | 12.9 | 11.9 | 10.8 |
| 2000 Hz | 20.2 | 19.3 | 18.3 | 17.4 | 16.4 | 15.5 |
| 4000 Hz | 19.3 | 18.5 | 17.6 | 16.8 | 16.0 | 15.2 |
| 8000 Hz | 17.3 | 16.6 | 15.9 | 15.2 | 14.6 | 13.9 |

<sup>1</sup>RETSPL for the ER-2 transducer (calibrated in an HA-1 coupler) was added to the model-estimated correction factors: 500 Hz = 6 dB, 1000 Hz = 0 dB, 2000 Hz = 2.5 dB, 4000 Hz = 0 dB, 8000 Hz = -3.5 Hz

**Figure 7.**
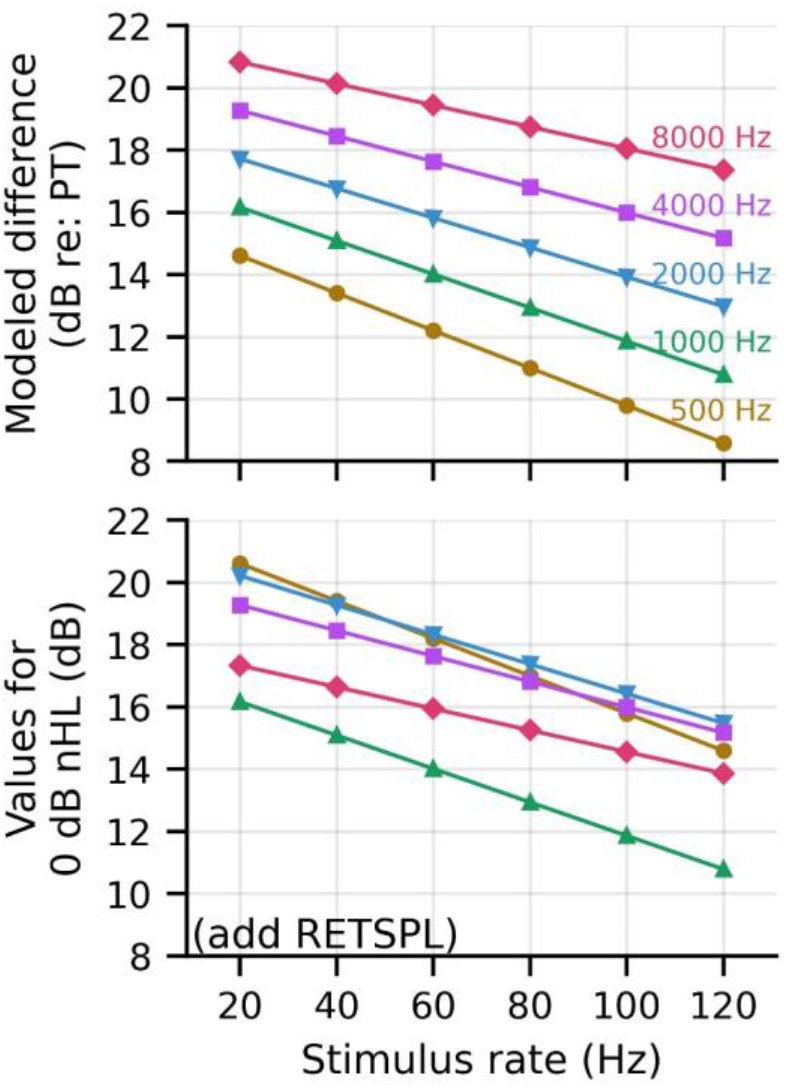
Correction factors for pABR stimuli. The modeled threshold differences, or correction factors, in (A) are added to the appropriate reference-equivalent threshold in SPL (RETSPL) for the transducer (in our study, ER-2 with HA-1 coupler) to give the normative values for audiometric zero (i.e., dB nHL) in (B). PT = pure tone

With these normative values, the 51 and 81 dB peSPL intensities used to evoke the pABR responses convert to a range from 30–35 and 60–65 dB dB nHL respectively for a 20 Hz stimulation rate to 36–40 and 66–70 dB nHL for 120 Hz rate. For the 40 Hz stimulation rate, 51 and 81 dB peSPL corresponded to 32–36 and 62–66 dB nHL respectively, with an average dB nHL across tone pip frequency of 33 and 63 dB nHL.

## DISCUSSION

Here, we describe how the pABR changes with stimulation rate and establish normative correction values for pABR levels based on perceptual thresholds. A wide range of rates yields robust responses in reasonable recording times for most subjects. For adults with normal hearing, the total recording time is limited by the broader component wave V of the waveforms for the low frequency tone pips. For some subjects with particularly broad responses, extending the analysis window improves response detection and the acquisition time necessary to reach SNR criterion. The perceptual thresholds to pABR stimuli subtly change with rate, giving a relatively similar set of correction factors to convert the level of the pABR stimuli from dB peSPL to nHL. Overall, the 40 Hz stimulation rate is the singular optimal rate, but in the clinic a range of rates may be useful for facilitating faster acquisition of elevated hearing thresholds across frequency.

Responses to pABR stimulation show adaptation to higher rates, both at a low and a high intensity. Even though the pABR uses random timing and simultaneous presentation of all 10 tone pips trains, wave V amplitude decreases with increasing frequency in a similar way to responses evoked by serially and randomly presented click stimuli (e.g., Burkard et al. 1990; Burkard & Hecox 1983; Chiappa et al. 1979; Don et al. 1977; Jiang et al. 2009; Burkard et al. 1996; Valderrama et al. 2014). The effect of rate on latency was not statistically significant here e, but the overall slight latency trend from 20 to 120 Hz were within the range of shifts previously reported for randomized click stimuli at higher rates (Burkard et al. 1996; Valderrama et al. 2014). Parallel presentation of randomized stimuli may also have contributed to differences in the effect of rate on latency. The benefit of higher rates is decreased noise, as variance reduces linearly with increasing number of stimuli. However, at some point adaptation limits this benefit by decreasing the response amplitude enough that SNR and response detection suffer. This trade-off can be seen in the estimates of recording time required to reach 0 dB SNR. The acquisition times improve (i.e., become faster) or remain similar for rates up to either 60 or 80 Hz – especially for the mid-to-high frequencies – and then lengthen again for the higher rates (Figure 3B) as the responses become smaller and broader (Figures 1, 4). When considering all tone pip frequencies at both intensities, the fastest recording times for most subjects is achieved with a 40 Hz stimulation rate – a rate that is consistently used in current clinical protocols (American Academy of Audiology 2012; BC Early Hearing Program 2012; Ontario Ministry of Children, Community and Social Services 2018). However, the trade-off between adaptation and SNR is subtle for the pABR stimuli, especially depending on the tone pip of interest. If focusing on low frequency tone pips then the 20–40 Hz rates are optimal, but for tone pips ≥ 2000 Hz, higher rates continue to improve acquisition times by a few minutes for most subjects (Figure 3). A few minutes represents valuable time when conducting diagnostic tests on infants, when the length of napping, and thus remaining testing time, is unknown. Furthermore, the effect of rate may not be as drastic for infants – amplitudes for click stimuli do not decrease as much for infants compared to adults, although they start off with a smaller amplitude for lower rates (Lasky 1997; Lasky 1984). Therefore, despite adaptation, a method that utilizes a combination of rates may speed up the time required to estimate hearing thresholds that may differ across frequencies, and will need to be tested in the clinic and with infants.

Acquisition times are reasonable for the pABR stimuli, and can be supported by extending the analysis window to better capture broader responses in some subjects. For most subjects, an analysis window of 10 ms adequately covers the wave V component (Figures 1, 5) and provides timely estimates to 0 dB SNR, with median total recording times for all 10 tone pips at most rates within 10 minutes for the lower intensity and within 4 minutes for a higher intensity (Figure 3). However, the recording time at a given level, at which the pABR yields responses for all frequencies in both ears, depends on the slowest response to emerge. For our adults with normal hearing, the broader 500 and 1000 Hz responses are the slowest – these two responses increase the median testing time for most rates from < 4 minutes for the lower intensity and < 2 minutes for the higher intensity to < 10 and < 4 minutes respectively. These are still acceptable times to simultaneously collect 10 responses and represent the substantial improvement in measurement time over serial measurement (Polonenko & Maddox 2019), but there are cases where the recording times are much longer for the low frequency tone pips. The “slowest quartile” of subjects have testing times that are < 10 minutes for 2–8 kHz tone pips but > 15 minutes for the low frequency tone pips (Figures 3, 5). In the cases where the low-frequency tone pip responses are visible, but the time for reaching the criterion SNR is taking longer than 6–9 minutes, there can be a significant speedup benefit of 5–16 minutes (corresponding to speedup ratios of 2–7) to using an extended analysis window that captures more of the broader response (Figure 4; Stapells & Oates 1997). This longer analysis window which is afforded by the random timing of the pABR and can be done for the same recording data, thereby not requiring extra recording runs/time. Next, the stimulation rate can be reduced to 20 Hz, which gives the most speedup advantage for detecting a response for 500 Hz at a lower intensity (Figure 5). This flexibility of using multiple analysis windows and stimulation rates gives another option in the toolkit that is easily implemented with the pABR.

Measurement time will also depend on hearing thresholds and implementation into a clinical setting. Time is limited by the 500 and 1000 Hz responses in the context of adults with normal hearing, but most hearing losses are more severe in the higher frequencies (e.g., Pittman & Stelmachowicz 2003). When obtaining responses at higher intensities, the threshold for the lower tone pip frequencies will either be established already or a response confirmed as present at the higher intensity. Then the time will be limited by the SNR of the higher tone pip frequency responses. Response amplitudes tend to be more linear for hearing loss and for high frequencies, because high frequency responses do not suffer the same blurring together of tone pips or adaptation (shallower amplitude-rate slopes at a level closer to threshold I Figure 2; smaller changes in recording time in Figure 3). Both of these suggest that there may still be time-saving advantages of using higher rates when searching for elevated thresholds at higher frequencies. At the lower intensity, 90% of subjects reached criterion for the 2–8 kHz tone pips within 5 minutes for the 80 Hz rate, compared to 7–9 minutes for rates < 80 Hz. Using faster rates for higher frequencies may save valuable time and allow more intensities to be tested within a recording session, giving a more complete exam. Of course the actual acquisition times will depend on the stopping criteria and implementation in the clinic. Here we used 0 dB SNR, but the reported minutes will scale if that threshold is changed (doubling, e.g., if it is increased to 3 dB SNR). We will next evaluate the pABR in a clinical setting and determine the time to find thresholds for a variety of degrees and shapes of hearing loss in adults who are able to sit through a complete session. Then, because the responses and acquisition times may differ with infants (Werner et al. 1993), we will finally test the method for evaluating hearing thresholds in infants.

Clinical implementation of the pABR also requires calibration of the pABR stimuli using perceptual thresholds. Perceptual thresholds to the brief pABR stimuli show some temporal integration with increasing rate (Figures 6, 7), but less than full integration. The energy increases nearly 6-fold – or 8 dB – from a stimulation rate of 20 to 120 Hz, but the differences in perceptual thresholds ranged from 3.5 dB for the 8000 Hz tone pip to 6 dB for the 500 Hz tone pip (Figure 7, Table 3). This minimal change in thresholds, and thus correction factors, across stimulation rates makes a multiple-rate paradigm easy to implement in an effort to obtain thresholds in the fastest recording time. For the optimal rate of 40 Hz, our normative values of 15.1–19.4 dB nHL for ER-2 earphones (Table 3, bottom) are lower than most reported values of 20–26 dB nHL for tone pips at similar stimulation rates (37.1–39.1 Hz, 41 Hz) using ER-3A inserts (Sharma et al. 2003; Stapells 2010; Stapells & Oates 1997). This holds even when we use our correction factors (Table 3, top) and convert to dB nHL using the RETSPLs for ER-3A inserts. This suggests that the pABR may require lower levels to obtain a response, but there are other variables that may contribute to these different norms. Primarily, our method adjusts for both temporal integration and the subject’s pure-tone threshold, whereas these other norms do not correct for thresholds < 15–20 dB HL (Sharma et al. 2003; Stapells 2010; Stapells & Oates 1997). Our values are more similar to the 17–21 dB nHL values obtained using the similar method of subtracting the pure-tone threshold from the threshold to the tone pip (Gorga et al. 1993). Other variables that contribute to some minor variation in norms are final step sizes (here we used 5 dB, others have used 2 dB), duration of the stimuli (like our study, most use 5 cycles, but some use 4 cycles), and calibration units for the tone pips (dB pSPL, peSPL, ppeSPL). We have provided the correction values in dB (Table 3, top) from which our normative dB nHL values were calculated, so that our values can be used by future studies by simply adding the RETSPLs for the transducers being used to the correction factors. We recommend using our correction factors because they directly compare our subjects’ thresholds to pure-tones and tone pips while using the same transducer, thereby accounting for temporal integration, hearing threshold re: 0 dB HL, and transducer. When using different transducers for future studies, however, it is important to note that the frequency responses of earphones differ, which may affect the morphology and amplitudes of the pABR responses to high-frequency tone pips in particular. For example, ER-3A inserts are commonly used in the clinical systems, which have a spectrum that decreases after 8 kHz. This is appropriate as most clinical diagnostic exams only test up to 4 kHz. But the pABR as implemented here tests up to 8 kHz, so we use the ER-2 inserts that have a flatter spectrum to 10 kHz. These differences in ER-2 and ER-3A transducer spectra have resulted in differences in ABRs to chirp stimuli (Elberling et al. 2012). Finally, these correction and normative values are for perception. Often, higher dB nHL levels are needed to evoke an electrophysiological response, and additional correction factors are needed to convert the physiological dB nHL level to estimated perceptual level (dB eHL). For example, for some systems, a normal hearing threshold of 25 dB eHL is established if an ABR response is present at 35–40 dB nHL for 500 Hz but 25 dB nHL for 4 kHz (American Academy of Audiology 2012; BC Early Hearing Program 2012; Ontario Ministry of Children, Community and Social Services 2018). Now that we have established the dB nHL corrections, our next step is to determine the relationship between dB nHL and dB eHL thresholds for a range of hearing loss severities and configurations.

For this study we focused on the wave V component of the pABR responses. However, additional earlier ABR and later middle-latency components are visible in the responses to pABR stimuli, particularly in at the higher intensity of 81 dB peSPL (Figures 1, 4). While not quantified herein, measuring the earlier waves may provide useful information for other clinical and research applications, such as evaluating cochlear synaptopathy (Bharadwaj et al. 2014; Bramhall et al. 2017; Liberman et al. 2016; Prendergast et al. 2017). Perhaps measuring responses at a low and high rate may reveal differences in synchronization at different stages of early auditory processing (Milloy et al. 2017). Often clicks of higher intensity (>100 dB peSPL) are used but even 81 dB peSPL yields robust responses with the pABR. Furthermore, results from our earlier work suggest that the pABR may be more place-specific at higher intensities than serially presented stimuli (Polonenko & Maddox 2019). Future studies can investigate the utility of the pABR for other applications than estimating hearing thresholds.

## CONCLUSIONS

The pABR evokes robust responses across a range of rates and intensities within reasonable recording times. The random timing affords extended analysis windows, allowing better estimates of noise in the pre-stimulus interval and faster detection of broader responses to low frequency tone pips or responses at lower intensities. A pABR method utilizing multiple stimulation may be useful in quickly estimating thresholds, particularly when thresholds are elevated at high but not low frequencies. We recommend that testing start with a stimulation rate of 40 Hz and a 10 ms analysis window. If responses are visible but longer times are needed to reach stopping criterion, then we suggest first increasing the analysis window to 30 ms and then change the rate as necessary. For calibration, we recommended using the correction factors established here and then adding the appropriate RETSPL for the specific transducer, so that the norms account for temporal integration, pure-tone threshold, and transducer. Finally, our future studies will evaluate the relationship between the ABR and perceptual thresholds as well as the measurement time in a clinical setting.

## Supporting information

Supplementary Digital Content 1

Supplementary Digital Content 2

## Abbreviations

ABR: – auditory brainstem response
ASSR: – auditory steady-state response
pABR: – parallel auditory brainstem response
dB peSPL: – decibels peak-equivalent (baseline-to-peak) sound pressure level
dB nHL: – decibels normal hearing level
PT: – pure tone
RETSPL: – reference equivalent threshold in sound pressure level

## ACKNOWLEDGMENTS

This work was supported by National Institute for Deafness and Other Communication Disorders (R00DC014288, R01DC017962) awarded to RKM. Portions of this article were presented at the 4^th^ International Hearing Loss Conference in Niagara-on-the-Lake, Ontario, Canada on May 6, 2019, and at the 2^nd^ Auditory EEG Signal Processing (AESoP) symposium in Leuven, Belgium on September 18, 2019. The article has been pre-published on BioRχiv (https://doi.org/10.1101/2021.05.13.444069). MJP designed and performed experiments, analyzed data and wrote the paper; RKM designed experiments, provided resources, discussed the results and implications and commented on the manuscript at all stages.

## DATA AVAILABILITY

EEG data will be made available in the EEG-BIDS format (Pernet et al. 2019) on Dryad (https://doi.org/10.5061/dryad.1c59zw3vm), as well as the stimulus files and python code necessary to derive the pABR responses. The behavioral data are deposited to the same Dryad repository.

## SUPPLEMENTAL DIGITAL CONTENT

**Supplemental Digital Content 1.** Table that describes details of the linear mixed effects models for wave V latency and amplitude that include the gender variable. pdf

**Supplemental Digital Content 2.** Figures that illustrate the CDFs for all stimulus rates and intensities using a 10 ms and 30 ms analysis window. pdf

