## Supplementary Digital Content 1 for "Optimizing Parameters for Using the Parallel Auditory Brainstem Response (pABR) to Quickly Estimate Hearing Thresholds"

### Supplemental Tables

***Supplemental Table 1.*** *Linear Mixed Effects Models for Wave V Latency and Amplitude*

| Fixed Effect | Estimate | SE | df | t | p | | Power (95% CI) |
|---|---|---|---|---|---|---|---|
| ***A. Latency formula:*** *latency ~ rate + intensity + logfreq + ear + rate:intensity + rate:logfreq + intensity:logfreq + logfreq:ear + rate:intensity:logfreq + gender + (1 \| subject)* | | | | | | | |
| Intercept [Left; Female] | 39.15 | 1.74 | 2260 | 22.45 | < 0.001 | *** | 1.00 (1.00, 1.00) |
| rate | -0.01 | 0.02 | 2250 | -0.27 | 0.784 | | 0.07 (0.05, 0.08) |
| intensity | -0.18 | 0.03 | 2250 | -6.94 | 0.000 | *** | 1.00 (1.00, 1.00) |
| logfreq | -8.11 | 0.52 | 2250 | -15.51 | < 0.001 | *** | 1.00 (1.00, 1.00) |
| ear[Right] | -0.97 | 0.34 | 2250 | -2.88 | 0.004 | ** | 0.82 (0.79, 0.84) |
| gender[Male] | 0.22 | 0.16 | 19 | 1.38 | 0.184 | | 0.29 (0.26, 0.32) |
| rate:intensity | 0.00 | 0.00 | 2250 | 1.74 | 0.082 | . | 0.42 (0.39, 0.46) |
| rate:logfreq | 0.00 | 0.01 | 2250 | 0.49 | 0.628 | | 0.06 (0.05, 0.08) |
| intensity:logfreq | 0.04 | 0.01 | 2250 | 5.18 | < 0.001 | *** | 1.00 (1.00, 1.00) |
| logfreq:ear[Right] | 0.26 | 0.10 | 2250 | 2.55 | 0.011 | * | 0.76 (0.73, 0.79) |
| rate:intensity:logfreq | 0.00 | 0.00 | 2250 | -1.69 | 0.091 | . | 0.40 (0.37, 0.43) |
| ***B. Amplitude formula:*** *amplitude ~ lograte + intensity + logfreq + ear + lograte:intensity + lograte:logfreq + intensity:logfreq + logfreq:ear + lograte:intensity:logfreq + gender + (1 \| subject)* | | | | | | | |
| Intercept [Left; Female] | 0.301 | 0.203 | 2266 | 1.48 | 0.139 | | 0.34 (0.31, 0.37) |
| lograte | -0.216 | 0.113 | 2261 | -1.91 | 0.056 | . | 0.50 (0.47, 0.54) |
| intensity | -0.006 | 0.003 | 2261 | -1.88 | 0.061 | . | 0.45 (0.42, 0.49) |
| logfreq | -0.069 | 0.061 | 2261 | -1.13 | 0.261 | | 0.21 (0.19, 0.24) |
| ear[Right] | -0.011 | 0.013 | 2261 | -0.86 | 0.392 | | 0.16 (0.13, 0.18) |
| gender[Male] | -0.025 | 0.012 | 19 | -2.05 | 0.054 | . | 0.54 (0.51, 0.57) |
| lograte:intensity | 0.002 | 0.002 | 2261 | 1.37 | 0.172 | | 0.28 (0.25, 0.31) |
| lograte:logfreq | 0.063 | 0.034 | 2261 | 1.85 | 0.065 | . | 0.48 (0.45, 0.52) |
| intensity:logfreq | 0.003 | 0.001 | 2261 | 3.68 | 0.000 | *** | 0.96 (0.94, 0.97) |
| logfreq:ear[Right] | 0.006 | 0.004 | 2261 | 1.62 | 0.105 | | 0.39 (0.36, 0.42) |
| lograte:intensity:logfreq | -0.001 | 0.001 | 2261 | -2.70 | 0.007 | ** | 0.78 (0.75, 0.80) |

*Note: CI = confidence interval; SE = standard error; logfreq = $\log_{10}$(frequency); lograte = $\log_{10}$(rate); . $p < 0.1$; * $p < 0.05$; ** $p < 0.01$; *** $p < 0.001$*
