## Supplementary Digital Content 2 for "Optimizing Parameters for Using the Parallel Auditory Brainstem Response (pABR) to Quickly Estimate Hearing Thresholds"

### Supplemental Figures

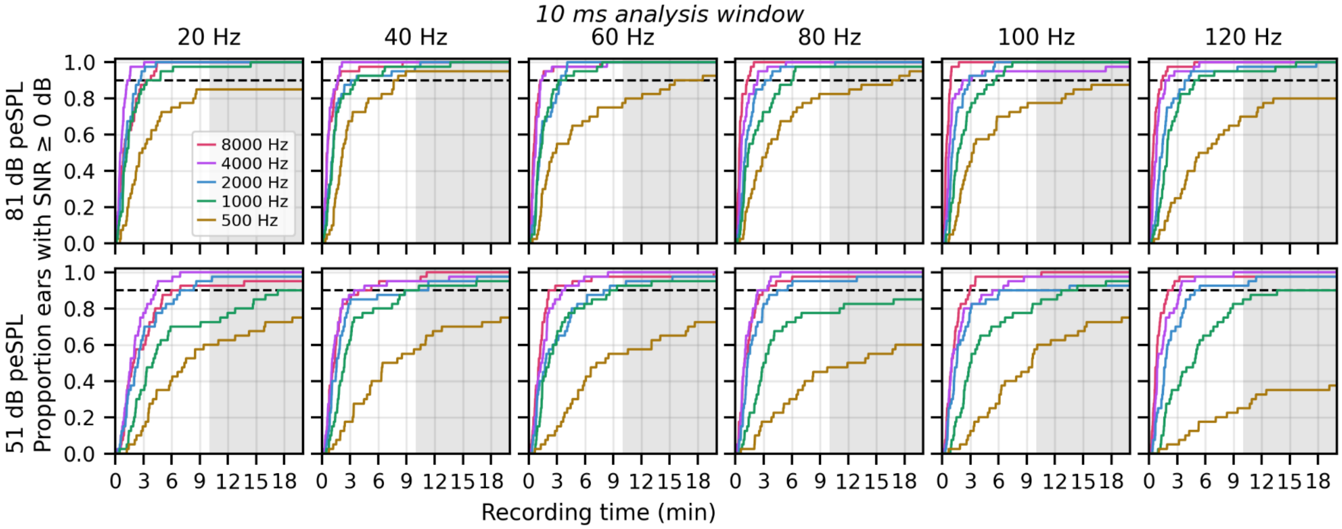


***Supplemental Figure 1.*** *Cumulative proportion of waveforms with ≥ 0 dB SNR as a function of recording time for a high (top) and low (bottom) presentation intensity for each tone pip frequency (indicated by line color) at each stimulation rate (columns). Shaded areas represent time estimations extrapolated past the 10-minute recording time. A 10 ms analysis window was used to calculate SNR.*

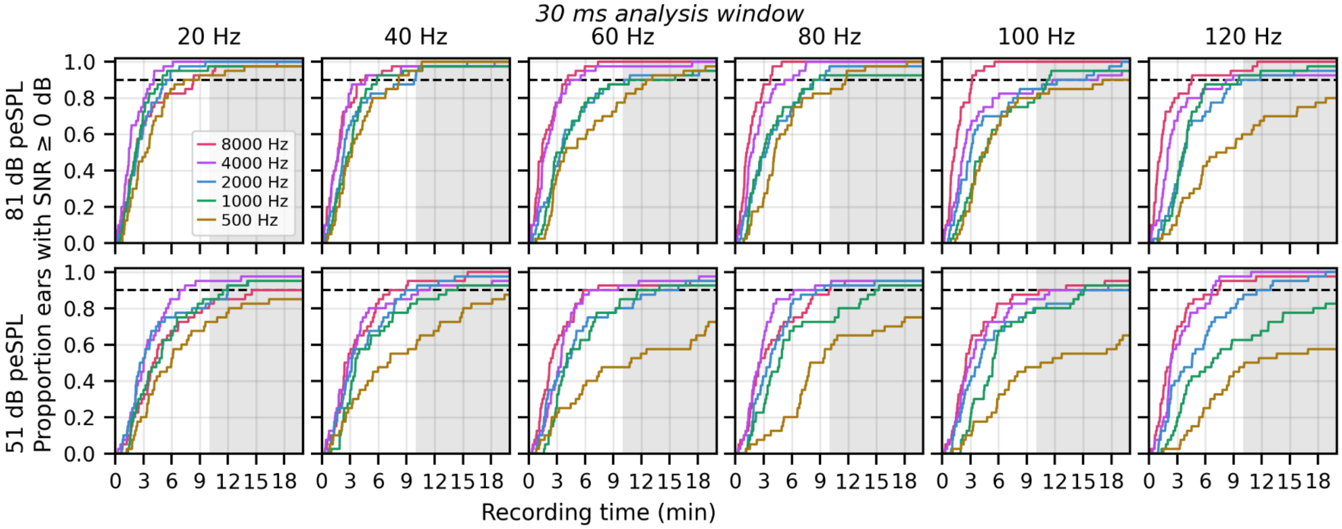


***Supplemental Figure 2.*** *Cumulative proportion of waveforms with ≥ 0 dB SNR as a function of recording time for a high (top) and low (bottom) presentation intensity for each tone pip frequency (indicated by line color) at each stimulation rate (columns). Shaded areas represent time estimations extrapolated past the 10-minute recording time. A 30 ms analysis window was used to calculate SNR.*
